# Compost bacteria as a promising new solution for degradation of diclofenac and related pharmaceuticals for water treatment processes

**DOI:** 10.1101/2025.07.09.663875

**Authors:** Francesca Demaria, Ramona Blattner, Chasper Puorger, Boris Kolvenbach, Mariana S Cretoiu, Philippe Corvini, Georg Lipps, Marcel Suleiman

## Abstract

Diclofenac, a widely used pharmaceutical, poses a significant environmental problem due to its persistence in aquatic systems and resistance to conventional degradation processes. Mesophilic microorganisms, commonly employed in wastewater treatment, often struggle to break down diclofenac, necessitating alternative approaches for its removal. In this study, we investigated thermophilic compost microorganisms and their ability to degrade diclofenac. Compost communities were cultivated for 20 weeks at 50°C in a membrane bioreactor, with a continuous supply of 2 mg/L diclofenac as the sole carbon source. After two weeks, the microbial community steadily enhanced its ability to remove diclofenac, achieving removal rates up to 60%. The consortium demonstrated flexibility in the degradation of further pollutants, namely sulfamethoxazole, paracetamol, and ciprofloxacin, with changes in their community structure depending on the substrates. In addition, thermophilic isolates *Chelatococcus* sp. strain D3 and *Mycobacterium* sp. strain D1 were characterized and demonstrated variation in the first reaction of transforming diclofenac, which is the crucial step in mineralization of this pollutant, resulting in either 4-hydroxy-diclofenac or diclofenac-lactam, respectively. Furthermore, *Chelotococcus* sp. strain D3 demonstrated the capability to catalyze the biotransformation of diclofenac into 4-hydroxydiclofenac in treated wastewater. Notably, this transformation was effectively carried out even at lower temperatures (25 °C and 37°C). These results show that the use of thermophilic consortia can be applied for efficient bioremediation in wastewater treatment plants, specifically for compounds that mesophilic organisms degrade poorly.

## Introduction

Water resources are in critical need of protection due to increasing contamination, overuse, and climate change, which threaten the availability and quality of this essential resource for ecosystems and human survival [1,2]. One prominent environmental concern is the widespread use of diclofenac, a non-steroidal anti-inflammatory drug (NSAID), which poses risks due to its persistence in the environment and potential toxicity [3–5]. Traditional wastewater treatment processes are often inadequate in completely removing diclofenac, leading to its accumulation in aquatic environments[6]. This persistence is primarily due to the chemical’s stability and resistance to biodegradation, allowing it to bypass conventional treatment barriers and enter natural ecosystems. Once in the environment, diclofenac can accumulate and adversely affect aquatic life [4,7], leading to ecological imbalances and potential human health risks through bioaccumulation in the food chain[8].

Biodegradation by microbial communities, along with activated carbon adsorption[9], ozonation[10], and advanced oxidation processes (AOPs)[11], is a widely used method for removing diclofenac from wastewater in treatment plants. However, while mesophilic bacteria have been reported to degrade diclofenac under certain conditions, their effectiveness is highly debated and often not consistently reproducible [12]. Diclofenac can still be detected in effluents from various WWTPs, thus the degradation rates *in situ* are generally low (WWTPs)[3]. This suggests that relying solely on mesophilic bacterial activity is inadequate for the effective biological removal of diclofenac from wastewater. The transformation of diclofenac to 4-hydroxy-diclofenac is recognized as the crucial step for biodegradation of diclofenac [13], and various studies showed that microbial cytochrome P450 monooxygenases can catalyze this transformation[14,15]. While diclofenac has a strong recalcitrant character, 4-hydroxy-diclofenac was shown to be easier degradable in various environmental compounds, indicating that 4-hydroxy-diclofenac appears to be the starting point for the efficient breakdown of diclofenac [13]. Besides 4-hydroxy-diclofenac, also Diclofenac-lactam (1-(2,6 dichlorophenyl)-2-indoline) was identified as a main TP for diclofenac in wastewater [16]. Furthermore, this metabolite was the main product of degradation when diclofenac was treated with UV/Se NPs/H2O2 nanoparticles[17], as well as in in water samples in which diclofenac was treated via acidic extraction [18]. While metabolic pathways for further degradation of 4-hydroxy-diclofenac were discussed [19], the further conversion of diclofenac-lactam remains unknown.

New biological systems need to be investigated to efficiently remove diclofenac from wastewater. We hypothesize that thermophilic compost bacteria could offer a promising solution for diclofenac degradation due to several compelling reasons: (i) Thermophiles are known for their ability to degrade recalcitrant substrates, since their enzyme machineries are extremely robust and could be more effective in breaking down complex pharmaceutical compounds [20–22]. (ii) Compost, which naturally hosts thermophilic bacteria, consists of chemical structures comparable to diclofenac, such as aromatics[23] and lignins [24], suggesting these bacteria may already possess the enzymatic pathways needed to degrade diclofenac. (iii) Moderate heat could enhance the bioavailability of diclofenac, making it more accessible for microbial degradation[25]. Thermophilic microorganisms have already been reported for the use in bioremediation and pollutant removal [21,26–28]. Exploring the potential of thermophilic compost bacteria for implementation in WWTP could lead to the development of more effective wastewater treatment strategies, significantly reducing the environmental impact of pharmaceutical pollutants like diclofenac and safeguarding aquatic ecosystems and human health.

## Materials and methods

### MBR performance and batch cultures

A laboratory-scale membrane bioreactor (MBR) was established with a total volume of 1 liter and filled with 500 mL of medium. Brunner mineral medium was used, consisting of the following components for 1 L: Na_2_HPO_4_ 2.44 g, KH_2_PO_4_ 1.52 g, (NH_4_)_2_SO_4_ 0.5 g, MgSO_4_ x 7 H_2_O 0.2 g, CaCl_2_ x 2 H_2_O 0.05 g and Trace element solution SL-4 10 ml. In addition, 2 mg/L sodium diclofenac was added as sole carbon source to the medium. After 15 weeks of operation, 0.05 g/L ammonium-acetate was added. Continuous aeration was provided using compressed air at a pressure of 0.5 bar and an oxygen concentration of 20 %. To ensure homogenization, a 3-cm stirrer operated at 400 rpm was employed. The MBRs featured a steel membrane holder equipped with two ultrafiltration membranes, each with a pore size of 0.08 µm, covering a total membrane area of 30 cm². The flow rate was maintained at 10 mL/h (hydraulic retention time (HRT) 50 hours), allowing the MBRs to operate continuously for a period of 20 weeks at a temperature of 50 °C.

The MBR was inoculated with a 60 °C compost sample taken from a composting site in Switzerland, by adding 10 g compost to the reactor. The MBR was sampled weekly by taking 2 ml culture: the supernatant was used for HPLC analysis, the pellet was used for DNA and for subsequent sequencing the V4 region of the 16S rRNA gene.

Batch cultures were performed using MBR consortia of week 20 for inoculation (1 % v/v) in 20 mL Erlenmeyer flasks, containing of Brunner medium as described above. As carbon source, 20 mg/L sodium diclofenac, ciprofloxacin, sulfamethoxazole, paracetamol, ibuprofen, and trimethoprim were used, respectively. Cultures were incubated for 6-20 days and sampled on day 6 for further microbial community analysis.

Isolates were gained by serial dilution of the MBR consortium, followed by plating on agar plates consisting of the mentioned medium as described above, containing 20 mg/L diclofenac, 5 ml/L glycerol and 50 mg/L acetate as carbon sources. Plates were incubated at 50 °C for three days, and grown colonies were screened in Erlenmeyer flasks containing 5 mL medium supplemented with 2 mg/L diclofenac at 50°C, 160 rpm. Two strains, which showed the most efficient removal capacity, were further analyzed.

One isolate, *Chelatococcus* sp. strain D3, was further evaluated in the previously described BMM medium, which was supplemented with 20 mg/L diclofenac and 50 mg/L acetate. The experiments were conducted at three different temperatures (25°C, 37°C, and 50°C), and additional tests were performed by introducing 20 mg/L each of sulfamethoxazole and trimethoprim as substrates. Further, diclofenac removal efficiency of the isolate *Chelatococcus* sp. strain D3 was tested in real treated wastewater taken from our in-house wastewater treatment plants (FHNW). The water parameters are shown in Table 1. The batch cultures was set up adding 20 mg/L of diclofenac and OD 0.1 of biomass. Samples were taken at the time of inoculation and after 48 hours and measured through HPLC.

**Table 1.**
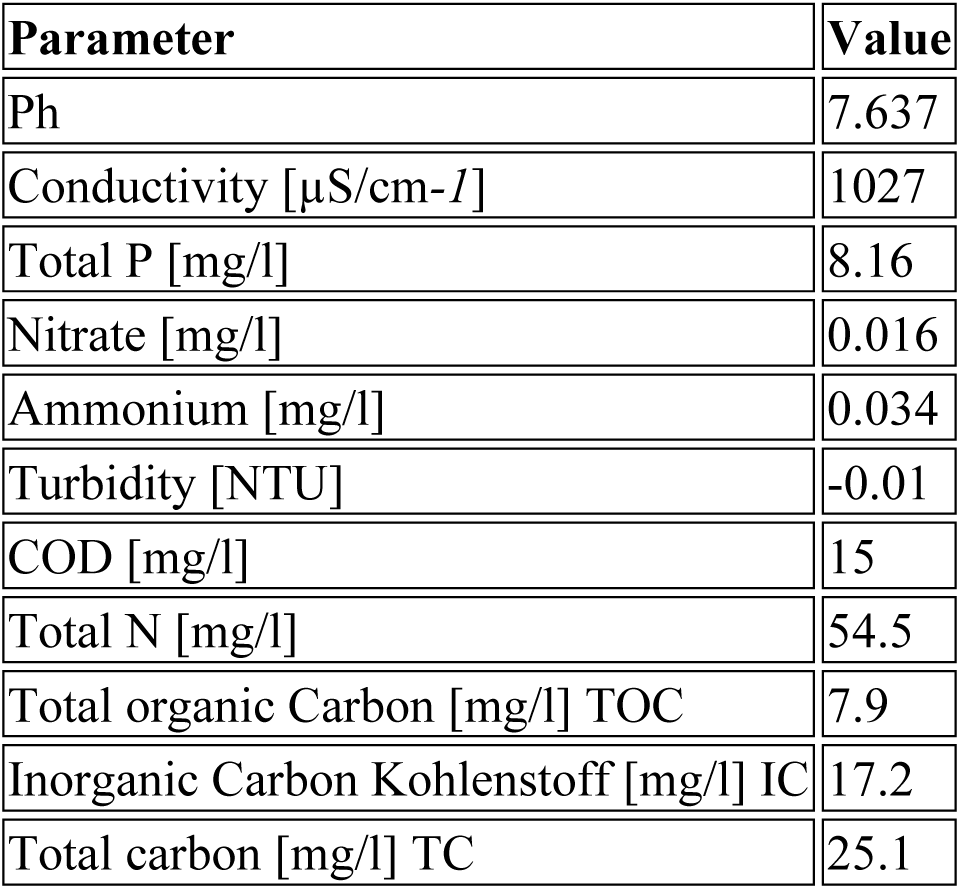
Characterisitcs of the treated wastewater used for inoculation with *Chelatococcus* sp. strain D3.

### DNA extraction, sequencing, and data analysis

DNA extraction was performed using the ZymoBIOMICS DNA Miniprep Kit (ZymoResearch), following the manufacturer’s protocol. The V4 region of the 16S rRNA gene was then amplified, and a DNA library was constructed using the Quick-16S™ Plus NGS Library Prep Kit (V4) from ZymoResearch. A 4 pM DNA library, spiked with 25% PhiX, was sequenced in-house using the Illumina MiSeq platform, adhering to the manufacturer’s guidelines. Sequencing data were processed based on primer sequences, quality, error rates, and chimeras using the ‘dada2’ R package [29]. The resulting sequence table was aligned with the SILVA ribosomal RNA database [30], version 138 (non-redundant dataset 99). A phyloseq object was created using the ‘phyloseq’ R package [31], incorporating the amplicon sequence variant (ASV) table, taxonomy table, and sample data. Prediction of metagenome functions was performed by PICRUSt2 [32]. Statistical analysis of microbial strain distribution was conducted by DESeq2 [33]. The phyloseq object, metadata, and detailed R analysis code are available on GitHub (https://github.com/Marcel29071989), and the raw sequencing data can be accessed at NCBI SRA under accession number SUB15191621.

### HPLC analysis

Diclofenac and its metabolites were separated on a ZORBAX RR StableBond C18 column using high-performance liquid chromatography (HPLC) (Agilent Technologies) by applying a flow rate of 0.7 mL/min with water and methanol as mobile phase. Diclofenac was detected using UV/VIS DAD detector. The mobile phase ratio started at 80:20 VV of, respectively, 0.1% formic acid in Millipore water (A) and methanol (B). The B gradient was programmed to transition from 20% to 95% over a span of 15 minutes, enabling the detection diclofenac at 13.8 minutes at a detection wavelength of 230 nm. A standard curve was generated for diclofenac (0.1 mg/L–100 mg/L).

### Mass spectrometry

Samples were thawed at room temperature, mixed and centrifuged for 5 min at 21,300 *g* (5425R, Eppendorf). For contaminant precipitation, 300 mL culture sample were mixed with 300 ml methanol and centrifuged for 5 min at 21300 *g*.

LC-MS/MS parameters: possible metabolites were identified from scans in the m/z range of 100-500 and compared with published degradation products. For quantification an LC-MS/MS method was set up using pure substances as reference. The chromatographic separation was done on an InfinityLab Poroshell 120 EC-C18 column (3.0 x 100 mm, 2.7 mm) connected to an Agilent 1100 HPLC system. The column was equilibrated in solvent A (5% methanol, 95% H_2_O with 0.1% formic acid) at a flow-rate of 0.5 ml/min. 2 ml of sample were injected and analytes were eluted with a linear gradient from 0% to 100% solvent B (95% (v/v) methanol, 5% H_2_O with 0.2% (v/v) formic acid) over 15 min. Analysis was performed with an Agilent 6470 QQQ MS equipped with an AJS-ESI ion source in positive or negative ion mode with a capillary voltage of 3500 V.

**Table 2.** Parameters of the MS method for detection of diclofenac and metabolites.

|  | m/z Precursor<br>ion | m/z Quantifier<br>(collision energy/eV) | m/z Qualifier<br>(collision energy/eV) | Ion mode |
| --- | --- | --- | --- | --- |
| Diclofenac | 296 | 214.1 (39) | 250 (13) | positive |
| 4'-hydroxydiclofenac | 310 | 266.2 (12) | 230.3 (8) | negative |
| 1-(2,6-Dichlorophenyl)-2-indolinone | 278 | 214 (33) | 208 (29) | positive |
| 2-hydroxyphenylacetic acid | 151 | 107.1 (12) | 106.4 (28) | negative |
| 2-aminophenylacetic acid | 152 | 134.4 (5) | 77.1 (46) | positive |

For quantification, standard curves of blank medium spiked with analyte concentrations between 2 and 2000 ng/ml were measured. Sample preparation procedures were the same as for the culture samples described above. Analyte contents in culture samples were determined using the obtained standard curves.

## Results

### Diclofenac degradation of the MBR community and possible microbial key players

Diclofenac (2 mg/L) removal was observed after three weeks of cultivation in the MBR, with 13% of the diclofenac being removed (Fig. 1a). From this point until Week 8, the removal efficiency increased steadily, reaching 58%. However, by Week 11, the removal capacity dropped to 9%. Starting at this time, ammonium acetate was added as a co-substrate alongside diclofenac. This addition improved the removal efficiency, which ranged between 33% and 58% from Week 15 to Week 20. No metabolites were detected in the effluent by mass spectrometry, except for small concentration of 4-hydroxy-diclofenac (>21 µg/L), indicating a complete mineralization of the removed diclofenac.

**Figure 1.**
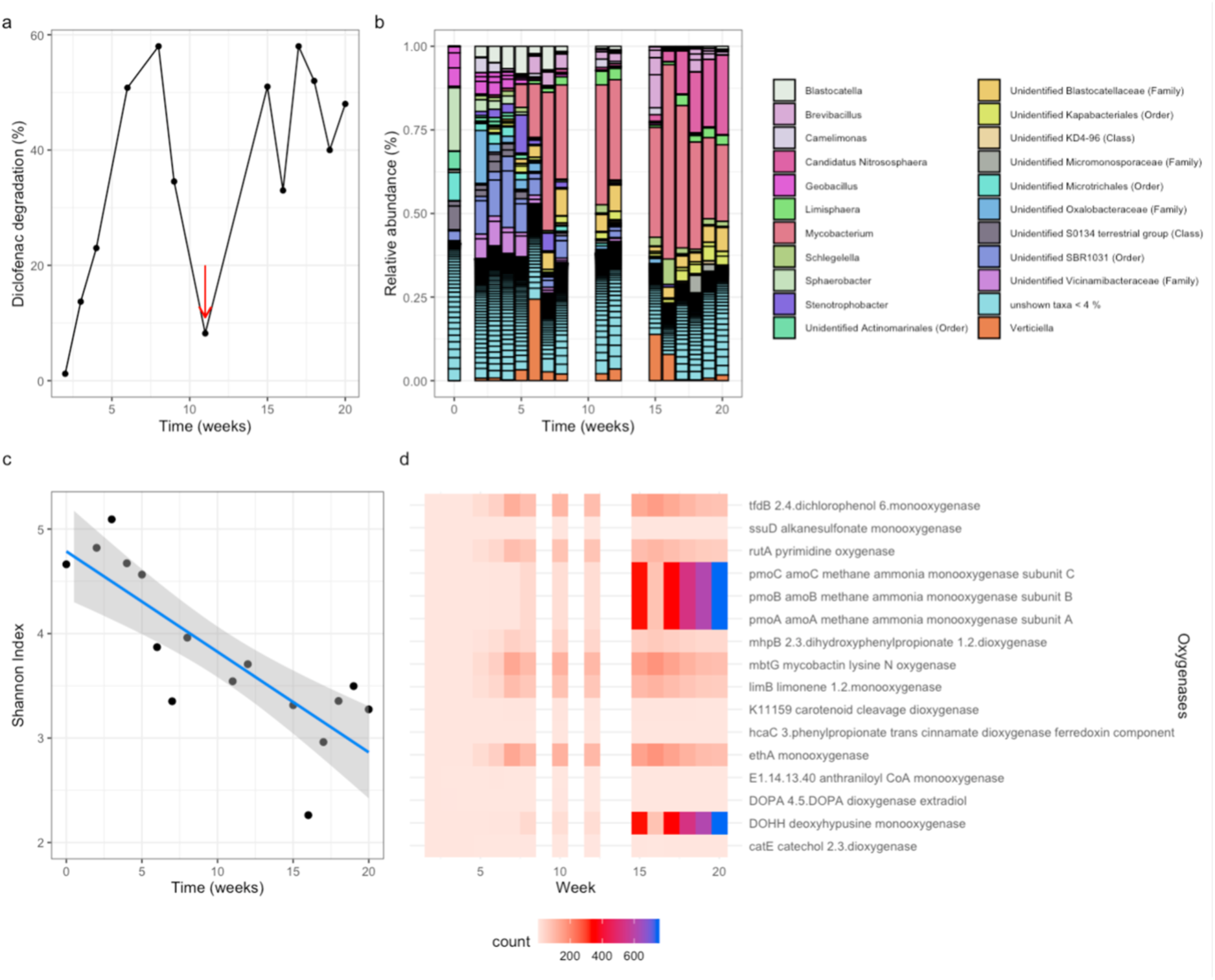
Performance and microbial community analysis of the diclofenac-degrading MBR. **(a)** Degradation (in %) of diclofenac within the MBR over time. Diclofenac concentration in the influent of the MBR was constantly 2 mg/L (Hydraulic retention time 50 h). The red arrow indicates the time point of addition of 50 mg/L ammonium-acetate within the influent. **(b)** Microbial community composition (relative abundance in %) within the MBR over time. Only ASVs with relative abundance > 4 % are identified for better visualization. **(c)** Shannon diversity index of the microbial community within the MBR. (d) Relative abundance of genomic oxygenases of the microbial community within the MBR. Oxygenases were initially selected based on their abundance (threshold 0.03 %) plotted based on the increase in count over time.

For each time point of diclofenac measurement, microbial community analysis was performed. The initial compost sample taken at 60 °C was primarily dominated by *Sphaerobacter thermophilus* (30.4%), followed by unidentified members of *Microtrichales* (13.5%) and *Gemmatimonadota* (11.4%), along with numerous unknown genera (Fig. 1B). After six weeks of incubation, a stable microbial community formed within the MBR, predominantly consisting of an unidentified genus from the *Alcaligenaceae* family, *Mycobacterium hassiacum*, and *Brevibacillus*. Subsequently, the relative abundance of *Mycobacterium hassiacum* increased significantly, reaching 64% by Week 16. Following the addition of ammonium citrate in Week 11, the ammonium oxidizer *Candidatus Nitrosophaera* also showed a marked increase, reaching 30% by Week 20. When the MBR reached its maximum degradation capacity in Week 17, the microbial community was composed of *Mycobacterium hassiacum* (42%), *Candidatus Nitrosophaera* (13 %), an unidentified member of *Blastocatellaceae* (3.5 %) and *Limisphaera* (3.2 %).

The diversity of the incubated microbial community within the MBR decreased over time, with a Shannon Diversity Index of 4.7 in the original compost sample, to a Shannon Diversity Index of 3.3 after week 20 of cultivation, indicating the microbial community became increasingly specialized on the compound diclofenac and its metabolites (Fig. 1 c).

Using PICRUSt2, the metabolic pathways of the microbial communities were analyzed, with a special focus on monooxygenases, that are known to play the crucial role to transform Diclofenac to 4-hydroxy-diclofenac. The most relative abundant oxygenases found in the community were analyzed regarding their initial value at week 0 (Fig. 1 d). Notably, the relative abundance of a methane ammonia monooxygenases increased during the 20-week incubation, as well as a DOHH deoxhypusine monooxygenase.

### Degradation of recalcitrant pharmaceuticals by the MBR consortium in batch cultures

As a next step, we tested if the adapted microbial community of the MBR is also able to degrade further recalcitrant pharmaceuticals in batch cultures, each incubated with a concentration of 20 mg/L (Fig. 2a). The HPLC results revealed that the communities were able to successfully remove diclofenac (74% in 20 days), Ciprofloxacin (45% in 20 days), Paracetamol (100% in 7 days), and Sulfamethoxazole (33% in 20 days). However, no removal was detected for Ibuprofen and Trimethoprim. Further, it was analyzed how the initial microbial community changed during cultivation in relation to the pollutant present in the batch culture (Fig. 2 b). Significant changes in the microbial community structures were observed after 10 days of incubation, with the extent of these alterations varying according to the specific pharmaceutical present (Fig. 2b). This shift is further highlighted in the NMDS analysis (Fig. 2c). The taxa *Mycobacterium*, unidentified *Blastocatellaceae*, and *Limisphaera* were among the most abundant in all cultures. Their relative abundances were influenced by the pharmaceutical present, with fluctuations observed depending on the compound.

**Figure 2.**
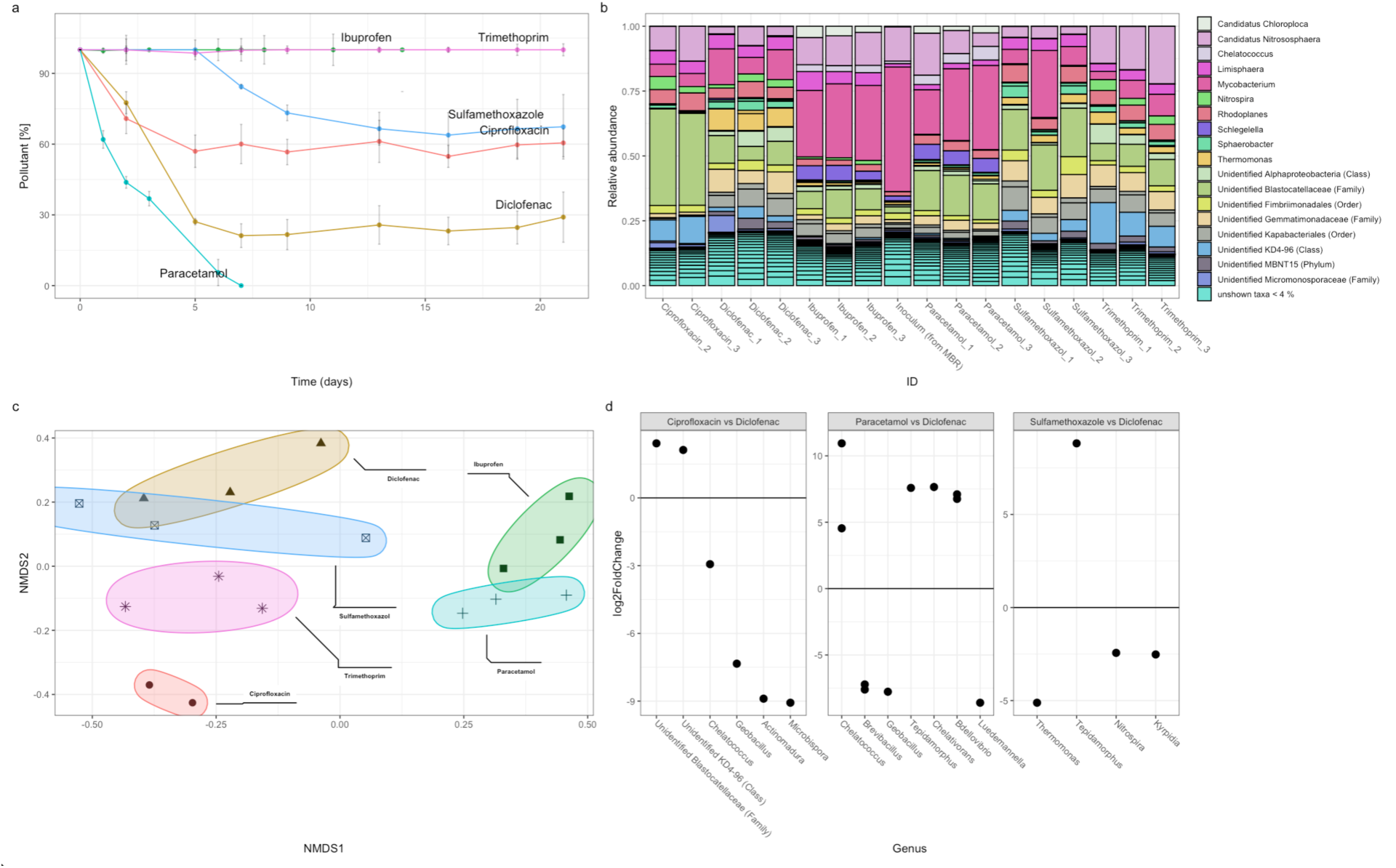
Performance and analysis of the MBR community within batch cultures facing various pharmaceuticals. **(a)** Concentration (in %) of Ciprofloxacin, Diclofenac, Ibuprofen, Paracetamol, Sulfamethoxazole and Trimethoprim within batch cultures inoculated with the adapted MBR community of week 20. Initial concentration of the pharmaceuticals was 5 mg/L, respectively **(b)** Microbial community composition (relative abundance in %) within the batch cultures on day 10 of cultivation. Just ASV > 0.04 are identified for better visualization. **(c)** NMDS analysis of the batch cultures. Stress is 0.11. (d) DESeq2 analysis of significant changes in the abundance of specific taxa between the batch cultures with different compounds. Error bars show ±SE (n = 3)

To analyze in detail which low-abundance microorganisms were significantly enriched in response to different substrates, we performed a DESeq2 analysis based on the relative abundance of taxa across treatments, using diclofenac as the reference condition (Fig. 2d). This approach also allowed us to detect trends in low-abundance taxa. In treatments containing ciprofloxacin, an unidentified candidate from the KD4-96 class and an unidentified member of the *Blastocatellaceae* family were significantly more abundant. In contrast, diclofenac treatments showed significantly higher abundance of the genera *Chelatococcus*, *Geobacillus*, *Actinomadura*, and *Microbispora*. When comparing microbial communities in paracetamol treatments to those in diclofenac treatments, *Chelatococcus* remained significantly more abundant, along with *Tepidamorphus*, *Chelativorans*, and *Bdellovibrio*. The genus *Tepidamorphus* was also significantly more enriched in treatments with sulfamethoxazole (SMX) compared to those with diclofenac.

These findings suggest that the microbial community adapted in the diclofenac MBR is capable of degrading a wide range of recalcitrant pharmaceuticals commonly found in wastewater. Due to the diverse microbial pool within the community, it demonstrates notable functional flexibility, enabling efficient responses to a variety of pharmaceutical contaminants.

### Isolates from the MBR capable of removing diclofenac

Thermophilic isolates were obtained by plating on diclofenac-containing agar plates through serial dilution of the thermophilic MBR inoculum, and grown colonies were screened for diclofenac removal at 50 °C. Two pure isolates showed the most potential removal activity: *Chelatococcus* sp. strain D3 (100 % identity to *Chelatococcus composti)* and *Mycobacterium* sp. strain *D1 (*100% identity to *Mycobacterium hassiacum)*. Interestingly, both strains transformed diclofenac into different metabolites. *Chelatococcus* sp. strain D3 transformed 33 % of 20 mg/L diclofenac into equimolar concentrations of 4-hydroxy-diclofenac within 48 hours (ammonium acetate present as co-substrate) (Fig. 3a). No growth of the cultures was monitored. *Mycobacterium hassiacum* transformed 100 % of 10 mg/L diclofenac into diclofenac-lactam within 8 days, while growth of the culture was observed under the tested conditions (Glycerol as co-substrate) (Fig. 3b). Both strains stopped the metabolization of diclofenac after the mentioned transformation under the tested conditions.

**Figure 3.**
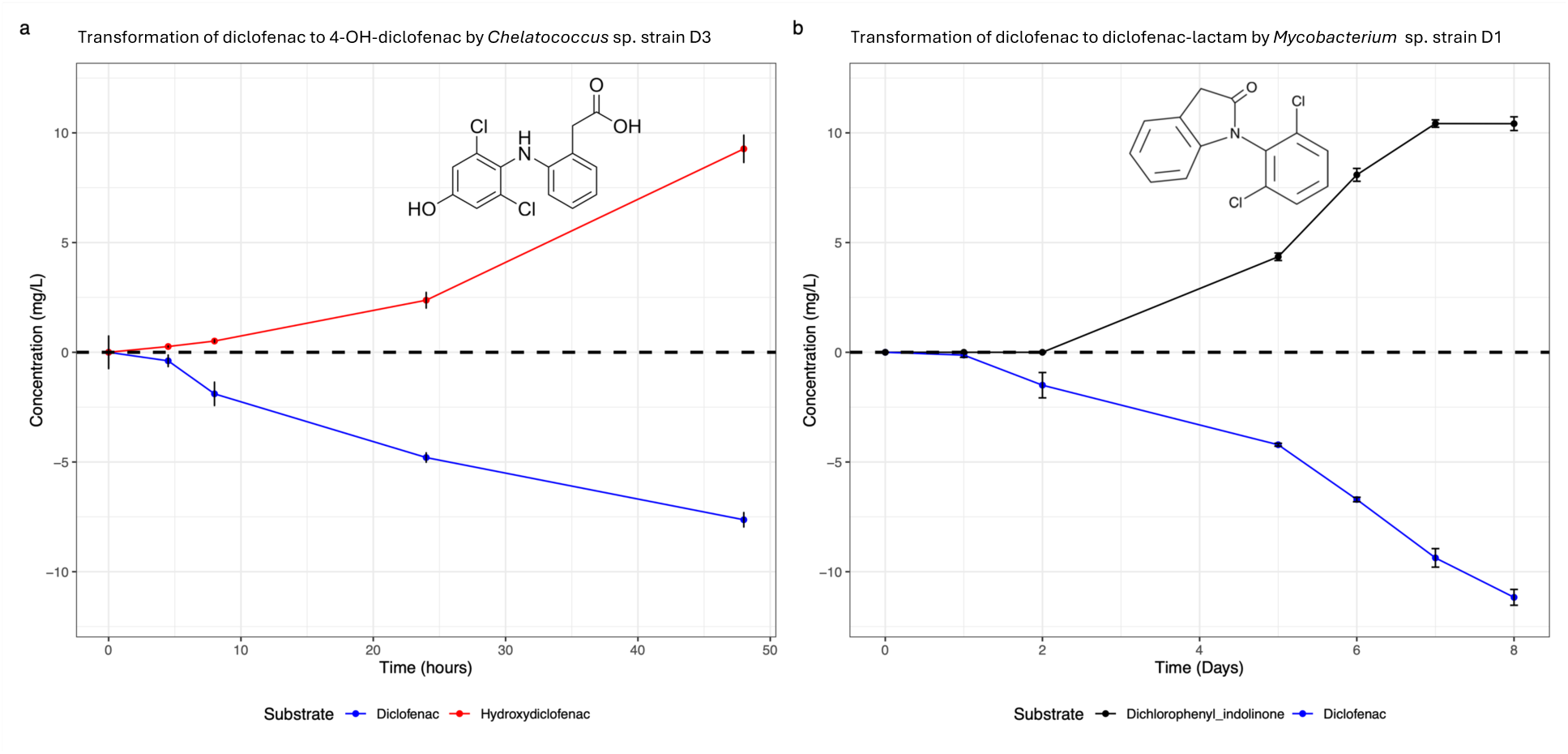
First reactions of transformation of diclofenac by 2 thermophilic compost bacterial strains at 50 °C. (a) Formation of 4-hydroxy-diclofenac by *Chelatococcus* sp. strain D3 within 48 hours (b) Formation of diclofenac-lactam by *Mycobacterium* sp. strain D1 after 8 days. Error bars show ±SE *(n = 3)*.

Due to its interesting formation of 4-hydroxy-diclofenac, the isolate *Chelatococcus* sp. strain D3 was further investigate on its performance at different temperatures at 25°C and 37 °C (and 50 °C for comparison) (Fig. 4a). At all temperatures, diclofenac was transformed to 4-hydroxy-diclofenac with slightly different efficiencies (20%, 25%, and 30% at 25°C, 37°C, and 50°C, respectively, within three days). Furthermore, *Chelatococcus* sp. strain D3 demonstrated the ability to remove diclofenac from real wastewater spiked with 20 mg/L of the compound (Fig. 4b). Impressively, it also degraded the antibiotics trimethoprim and sulfamethoxazole at an elevated temperature of 50 °C (Fig. 4c). These results highlight *Chelatococcus* sp. strain D3 as a highly promising candidate for the removal of diverse pharmaceutical contaminants from wastewater.

**Fig. 4.**
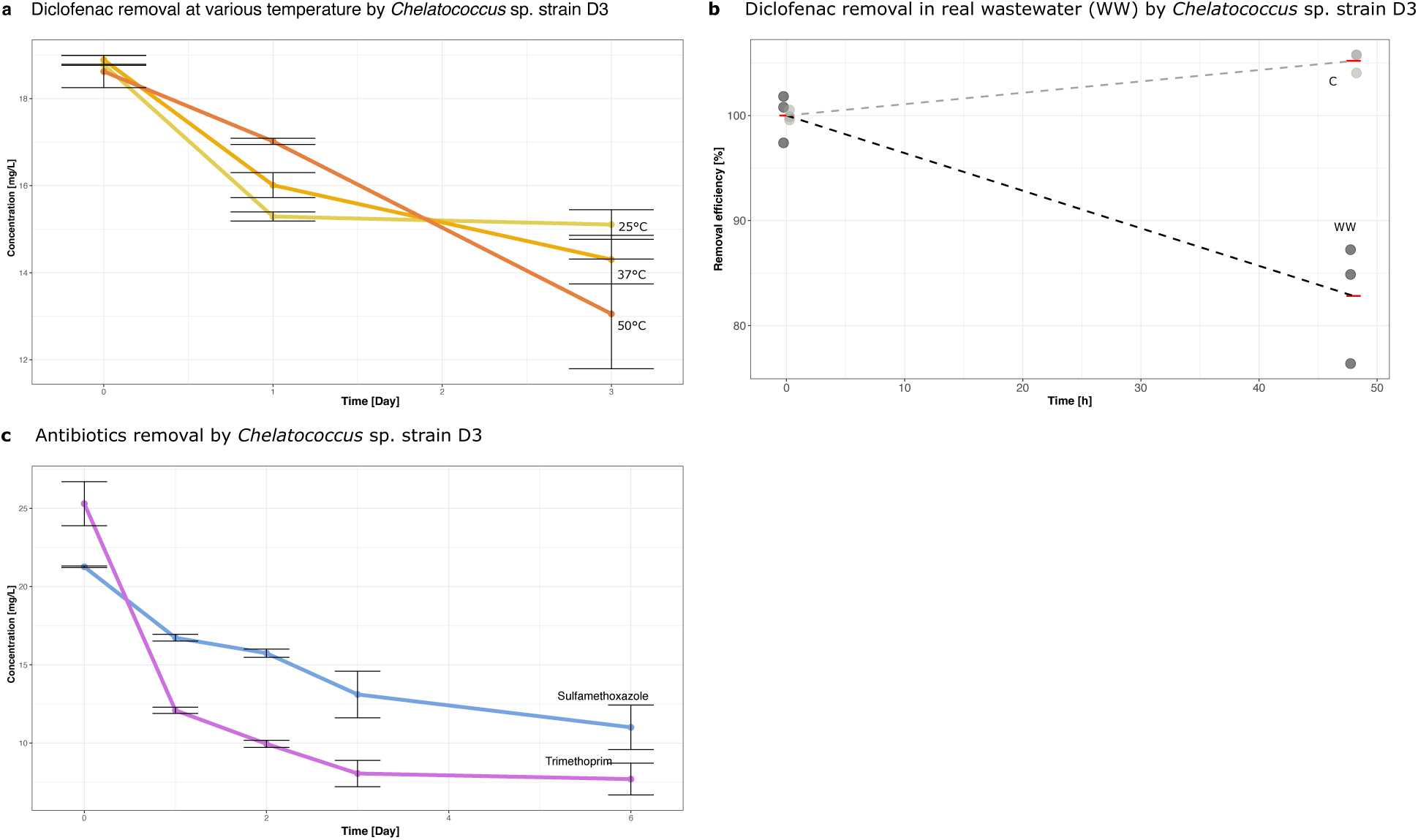
Characteristics of the degradation of pharmaceuticals by *Chelatococcus* sp. strain D3. (a) Diclofenac removal efficiency by *Chelatococcus* sp. strain D3 cultivated at different temperatures (25°C, 37°C, 50°C). Error bars show ±SE *(n = 3).* (b) Removal efficiency in % of diclofenac by *Chelatococcus* sp. strain D3 in real wastewater (WW) spiked-in with 20 mg/L of diclofenac at 50 °C. Abiotic controls represent batch cultures without addition of *Chelatococcus* sp. strain D3. Error bars show ±SE *(n = 3).* (c) Removal efficiency of the antibiotics sulfamethoxazole and trimethoprim by *Chelatococcus* sp. strain D3 at 50 °C. Error bars show ±SE *(n = 3)*.

## Discussion

The rising consumption of diclofenac and the low removal efficiency in wastewater treatment plants have led to the presence of this emerging contaminant in various environmental compartments, particularly in water. Diclofenac remains alarmingly detectable in the effluents of numerous wastewater treatment plants, defying attempts to remove it effectively[5,6]. Furthermore, this persistent contaminant has also been found in lakes and rivers, highlighting its widespread and concerning presence in our water sources [34]. This persistence is primarily due to the chemical’s stability and resistance to biodegradation, allowing it to bypass conventional treatment barriers and enter natural ecosystems. Given the persistent presence of diclofenac in wastewater effluents and aquatic environments, it is crucial to explore innovative biological solutions for its degradation. Our planet hosts a diverse array of microorganisms with complex metabolic pathways that could provide effective strategies for addressing this environmental challenge [35]. The study revealed that thermophilic microbial communities from compost can remove diclofenac, which shows new potential for facing the challenge of the tackling of this persistent compound. Moreover, the community established after 20 weeks within the MBR was also capable of removing the antibiotics sulfamethoxazole and ciprofloxacin, as well as the pain killer paracetamol, which are all present and persistent in wastewater [36–38]. This broad substrate range underscores the potential for using these microbial consortia in systems that treat complex pharmaceutical mixtures often found in wastewater effluents. The observation of community structure shifts based on the available pharmaceutical substrate suggests a dynamic response to environmental pressures [39,40], further supporting the adaptability and resilience of these microbial communities in real-world wastewater treatment conditions.

Addition of ammonium acetate as a co-substrate significantly improved the removal efficiency in later stages of the MBR operations. Therefore, it is very likely that diclofenac is broken down through co-metabolism[41], which is an often-observed phenomena for degradation of complex pharmaceuticals [42,43]. It is likely that the first weeks in the MBR operated without acetate, as natural substances from the compost could have served as co-substrates. Furthermore, the absence of diclofenac metabolites in the effluent suggests that degradation processes are taking place, and that adsorption can be ruled out as a contributing factor. This is also underlined by the fact that isolated strains could start the transformation reactions of diclofenac. The archaeon *Candidatus Nitrosophaera* occurred in the MBR after ammonium was present in the MBR, which oxidized the compound to nitrite. This may be advantageous in real-world scenarios since ammonium is highly present in municipal wastewater.

Future research should focus on scaling up these findings, optimizing operational conditions, and further investigating the enzyme systems involved in diclofenac degradation to maximize efficiency. Given the widespread occurrence of diclofenac in the environment, the development of robust and efficient treatment systems for its removal is essential to safeguard water quality and aquatic ecosystems.

A major limitation of this thermophilic wastewater treatment technology is the energy required to heat the system to 50 °C. However, processes with such moderate temperatures can be successfully carried out by utilizing district heating [44]. Furthermore, the isolate *Chelatococcus* sp. D3 was also able to degrade diclofenac at 25°C and 37 °C, making it a promising candidate for water treatment. Similarly, *Mycobacterium hassiacum* has been described in the literature to grow under mesophilic conditions, such as 35-40 °C [30]. This suggests that it may be possible to operate or adapt the compost community within the MBR to slightly lower temperatures, potentially reducing energy costs and improving environmental sustainability. On the other hand, the higher temperature might be necessary to enhance the bioavailability of diclofenac. The isolation of *Mycobacterium* sp. D1 and *Chelatococcus* sp. D3 is particularly interesting, as they represent both a highly abundant and a low-abundance member of the microbial community. *Mycobacterium* was found to be highly abundant, constituting more than 40% of the community, whereas *Chelatococcus* accounted for less than 4%. However, the importance of *Chelatococcus* becomes evident in the DESeq2 analysis of batch cultures grown with different substrates. This analysis revealed that *Chelatococcus* was significantly more abundant in cultures exposed to diclofenac (and paracetamol), suggesting a specific role in the degradation of these pharmaceuticals despite its overall lower abundance in the microbial community. Moreover, *Chelatococcus* was tested for the degradation of two antibiotics, sulfamethoxazole and trimethoprim. The degradation occurred in a fast fashion within 6 days, demonstrating its potential on bioremediation of further pharmaceuticals. This finding demonstrates that low-abundant bacteria can play a crucial role for the function of ecosystems [45]. Even though they make up a small fraction of the microbial population, their specialized metabolic capabilities can significantly influence the overall biodegradation process in complex microbial ecosystems. Further, the finding that both strains produced different metabolites from diclofenac highlights the diversity of metabolic reactions involved in its transformation. While the complete pathway for 4-hydroxy-diclofenac removal has been described in the literature, the metabolism via diclofenac-lactam remains only partially understood. Since diclofenac-lactam is not present in the MBR effluent of this study, it suggests that further transformation occurs beyond this activation step by using the full potential of a whole microbial community within the MBR. On the other hand, diclofenac-lactam has been identified as a major metabolite in wastewater [19], which could indicate that this pathway leads to more persistent compounds that are harder to degrade. This finding raises concerns about the efficiency of wastewater treatment processes in fully breaking down diclofenac and its transformation products. Further research is needed to clarify whether diclofenac-lactam represents a dead-end metabolite or if additional degradation steps exist.

In conclusion, this study highlights the potential of thermophilic compost-derived microbial communities to remove recalcitrant pharmaceuticals from wastewater treatment systems. The adaptability and metabolic versatility of these communities present a promising avenue for developing more effective strategies to mitigate pharmaceutical pollution in aquatic environments. Thermophilic membrane bioreactors (MBRs) may offer significant advantages for the advanced polishing of pharmaceutical-laden wastewaters, particularly those originating from industrial production and hospital effluents. Notably, Chelatococcus sp. strain D3 appears to be a particularly promising candidate for bioaugmentation in wastewater treatment plants (WWTPs), as it catalyzes the critical transformation of diclofenac to 4-hydroxy-diclofenac. This strain was also active at lower temperatures and demonstrated additional degradative capacity toward other recalcitrant pollutants, underscoring its potential role in enhancing the efficiency and robustness of biological wastewater treatment processes.

## Acknowledgements

Not applicable.

## Data availability

All datasets and metadata are available on the GitHub repository Marcel 29071989 (https://github.com/Marcel29071989/), and the raw sequencing data can be found on NCBI SRA archive under ID SUB15191621.

## Code availability

R Studio code for the analysis and plotting of figures for the manuscript and supplementary information is available at https://github.com/Marcel29071989

## Ethics approval and consent to participate

Not applicable.

## Consent for publication

Not applicable.

